# The field of view available to the ventral occipito-temporal reading circuitry

**DOI:** 10.1101/069369

**Authors:** Rosemary Le, Nathan Witthoft, Michal Ben-Shachar, Brian Wandell

**Author notes:** ^*^These authors contributed equally to this work (Rosemary Le, Nathan Witthoft). Contact: Rosemary Le.

## Abstract

Skilled reading requires rapidly recognizing letters and word forms; people learn this skill best for words presented in the central visual field. Measurements over the last decade have shown that when children learn to read, responses within ventral occipito-temporal cortex (VOT) become increasingly selective to word forms. We call these regions the VOT reading circuitry (VOTRC). The portion of the visual field that evokes a response in the VOTRC is called the *field of view (FOV)*. We measured the FOV of the VOTRC and found that it is a small subset of the entire field of view available to the human visual system. For the typical subject, the FOV of the VOTRC in each hemisphere is contralaterally and foveally biased. The FOV of the left VOTRC extends ~9° into the right visual field and ~4° into the left visual field along the horizontal meridian. The FOV of the right VOTRC is roughly mirror symmetric to that of the left VOTRC. The size and shape of the FOV covers the region of the visual field that contains relevant information for reading English. It may be that the size and shape of the FOV, which varies between subjects, will prove useful in predicting behavioral aspects of reading.

## Introduction

Learning to identify letters and words rapidly and reliably requires extensive visuospatial training. After training, the ability to rapidly recognize words varies across the visual field. Letters and words are recognized most efficiently in the central visual field; even when letter size is appropriately scaled, reading in the periphery remains slow (Chung, Mansfield, & Legge, 1998). With eccentric viewing, word recognition is faster for text presented in the lower visual field than in the left or right visual field (Petre, Hazel, Fine, & Rubin, 2000).

Neuroimaging and intracranial electrode measurements also provide evidence that regions in ventral occipitotemporal (VOT) cortex are part of the neural circuitry that subserves reading (Dehaene & Cohen, 2011; Kravitz, Vinson, & Baker, 2008a; Price & Devlin, 2011; A. M. Rauschecker, Bowen, Parvizi, & Wandell, 2012; Brian A. Wandell, Rauschecker, & Yeatman, 2012). These regions are involved in the perception of visually presented word forms and are reproducibly activated in fMRI experiments across individuals and orthographies (Baker et al., 2007; Bolger, Perfetti, & Schneider, 2005; Glezer & Riesenhuber, 2013; Krafnick et al., 2016; Martin, Schurz, Kronbichler, & Richlan, 2015). We analyze the regions within VOT that are more responsive to visually-presented words than to other categories of visual stimuli, and we refer to these regions as the VOT reading circuitry (VOTRC).

Yu et al. used neuroimaging and stimuli at different temporal frequencies to investigate why word identification is better in the fovea than the periphery (Yu, Jiang, Legge, & He, 2015). We further analyze the role of visual field in reading by measuring the portion of the visual field where the presentation of stimuli evokes responses in the VOTRC. We refer to this part of the visual field as the region’s field of view (FOV), and we measure it using population receptive field (pRF) methods (Dumoulin & Wandell, 2008; K. N. Kay, Winawer, Mezer, & Wandell, 2013).

The analyses described here quantify the FOV in the VOTRC in typical adult readers in a range of conditions. First, we assess the shape and position of the FOV in the VOTRC within the left hemisphere. We show that the FOV can be reliably measured and differs between subjects. The VOTRC responds to letters most strongly, but it also responds to simple contrast patterns. We find that the FOV of the same VOTRC voxels increases when we measure with checkerboards rather than words. We further analyze the responses in the VOTRC of the right hemisphere.

Given that responses in the VOTRC change with reading development (Michal Ben-Shachar, Dougherty, Deutsch, & Wandell, 2011a; Brem et al., 2006; Dehaene et al., 2010) it is likely that the visual information relayed to the VOTRC influences reading performance. The signals in this region may serve as a bottleneck, or may influence the way a subject reads (e.g., their eye movement pattern). The FOV measurements we describe can serve as a foundation for future analyses of reading in clinical and developmental populations.

## Methods

### Subjects

Twenty right-handed subjects (12 females, median age 24 y, range 20 - 35y) participated in the study, which was approved by the Institutional Review Board at Stanford University. All subjects gave informed consent, were right-handed native English speakers, and had normal or corrected-to-normal vision with no reported history of learning disabilities.

### Displays

Stimuli were presented with two types of displays. Retinotopic-mapping stimuli were presented with an Eiki LC-WUL100L projector. The Eiki projector has a native resolution of 1920 × 1200 pixels and 10-bit color resolution. Localizer stimuli were presented with either the Eiki projector or a 47″ LCD display. The 47″ LCD display has a native resolution of 1920 × 1080 and 8-bit color resolution. Both displays were viewed through a mirror placed ~5cm (projector) ~15cm (LCD display) from the eyes. Stimuli projected using the LCD display had a vertical extent of 12°, while stimuli presented with the Eiki projector had a vertical extent of 32°.

### MR Acquisitions

Data were acquired at the Stanford Center for Cognitive and Neurobiological Imaging using a 3T General Electric MR 750 scanner with either a Nova 16- or 32-channel head coil. The Nova 16-channel is similar to the Nova 32-channel coil, but with the front half of the coil removed to allow for an unobstructed field of view. Head motion was minimized by padding around the head.

#### Anatomical

Anatomical data were acquired with the 32-channel coil using a 3D Fast SPGR scan (166 sagittal slices, resolution 0.9375 × 0.9375 × 1mm). For each subject, 1-3 anatomical volumes were acquired and then averaged. Data were resampled to 1mm isotropic voxels and aligned to the anterior commissure - posterior commissure (AC-PC) plane using an affine transformation.

#### Functional

Functional data for pRF mapping and the large-field localizer (see below) were acquired with the 16-channel coil. 36 slices covering occipitotemporal cortex were defined: 2.5mm isotropic voxels, TR 2000ms, TE 29ms, flip angle 77°, field-of-view 200x200 mm. Functional data for the small-field localizer (see below) were acquired with the 32-channel coil. 48 slices covering occipitotemporal cortex were defined: 2.4mm isotropic voxels, TR 1000ms, TE 30ms, flip angle 62°, field-of-view 192 × 192mm. An in-plane anatomical image that matched the functional slice prescription was acquired before each set of functional runs. These images were used to align the functional data to the anatomical volume data.

### MR Data Analysis

#### Preprocessing

MR data analyses relied on the open-source code in vistalab (https://github.com/vistalab/vistasoft). The basic pre-processing steps included estimation and removal of motion artifacts, and registration of the functional data to the high resolution anatomical images. Motion artifacts within and across runs were corrected using an affine transformation of each volume in a session to the first volume of the first run. In all subjects, head movement was less than 1 voxel (in most cases less than 0.4 voxel). The first 6 time frames of each functional run were discarded. Baseline drifts were removed from the time series by high-pass temporal filtering.

The inplane anatomical image was aligned to the average whole brain T1-weighted anatomical image by calculating a rigid body transformation that maximized the mutual information between the inplane anatomy and the resliced volume anatomy. These alignment parameters were then used to align the functional data to the anatomical data.

#### Defining the VOT reading circuitry (VOTRC)

We defined the VOTRC using a combination of functional and anatomical constraints. One of two localizers was used to choose voxels based on their functional selectivity: a small-field or a large-field localizer. In the small-field localizer, stimuli consisted of a single item (pseudoword, body, face, place, object, number) overlaid on a phase-scrambled background (Figure 1A). Each phase scrambled background was generated from an image selected randomly from the entire set of images. The background image subtended a 12° × 12° portion of the visual field. For the word category, each word superimposed on the background spanned approximately 3° × 8°. We call this the small-field localizer because the category of interest was only present in the more central part of the visual field. The small-field localizer was used to identify the VOTRC in Subjects 1-12.

In the large-field localizer, a 30° × 30° portion of the visual field was tiled with either words, faces, or a phase-scrambled object (Figure 1B). The purpose of this localizer was to present words to both central and peripheral locations in the visual field in order to avoid biasing voxel selection to only voxels responsive to the central visual field. English words were presented rather than pseudowords (both words and pseudowords are well known to evoke strong responses in the VOTRC, (Dehaene, Le Clec’H, Poline, Le Bihan, & Cohen, 2002). The words were 1-6 letters long, had a minimum frequency of 200 per million, were displayed in Helvetica font, and the lowercase ‘x’ character had a vertical size of 1.3°. For each of the face images, 16 faces were chosen at random from a database of 144 faces and arranged in a 4×4 configuration. Each face had a vertical size of approximately 6°. In this localizer, participants were only presented with word, face, and phase-scrambled object categories. The large-field localizer was used to identify the VOTRC in Subjects 1, 13-20.

In both localizers, stimuli were presented in blocks of 8 images from the same category at a rate of 2 Hz. Each category was presented 6 times per run, and 3 runs were collected per subject. The categorical stimuli for both localizers were taken from the fLoc functional localizer package ((Stigliani, Weiner, & Grill-Spector, 2015), http://vpnl.stanford.edu/fLoc/). Participants fixated a small red dot at the center of the screen. Zero, one, or two phase-scrambled images appeared in a block with probabilities of 0.25, 0.5, and 0.25, respectively. Participants pressed a button when a phase-scrambled image appeared.

The boundaries of the VOT are the inferior temporal sulcus (lateral), the collateral sulcus (medial), hV4 (posterior), and an imaginary horizontal line drawn from the collateral sulcus to the inferior temporal sulcus, starting at the point where the parieto-occipital sulcus would meet the collateral sulcus (anterior). In most subjects, hV4 is found in the posterior transverse collateral sulcus (N. Witthoft et al., 2014). This region is illustrated in Figure 1C. The selective voxels within this region in each hemisphere are defined as the left or right VOTRC. Selective voxels are more responsive to words than to other visual categories (t-test, p<0.001, uncorrected), but not necessarily contiguous. The VOTRC falls in or near the occipital temporal sulcus (Figure 3A) in both hemispheres. The VOTRC as defined for all subjects is shown in Supplementary Figure 1.

**Figure 1.**
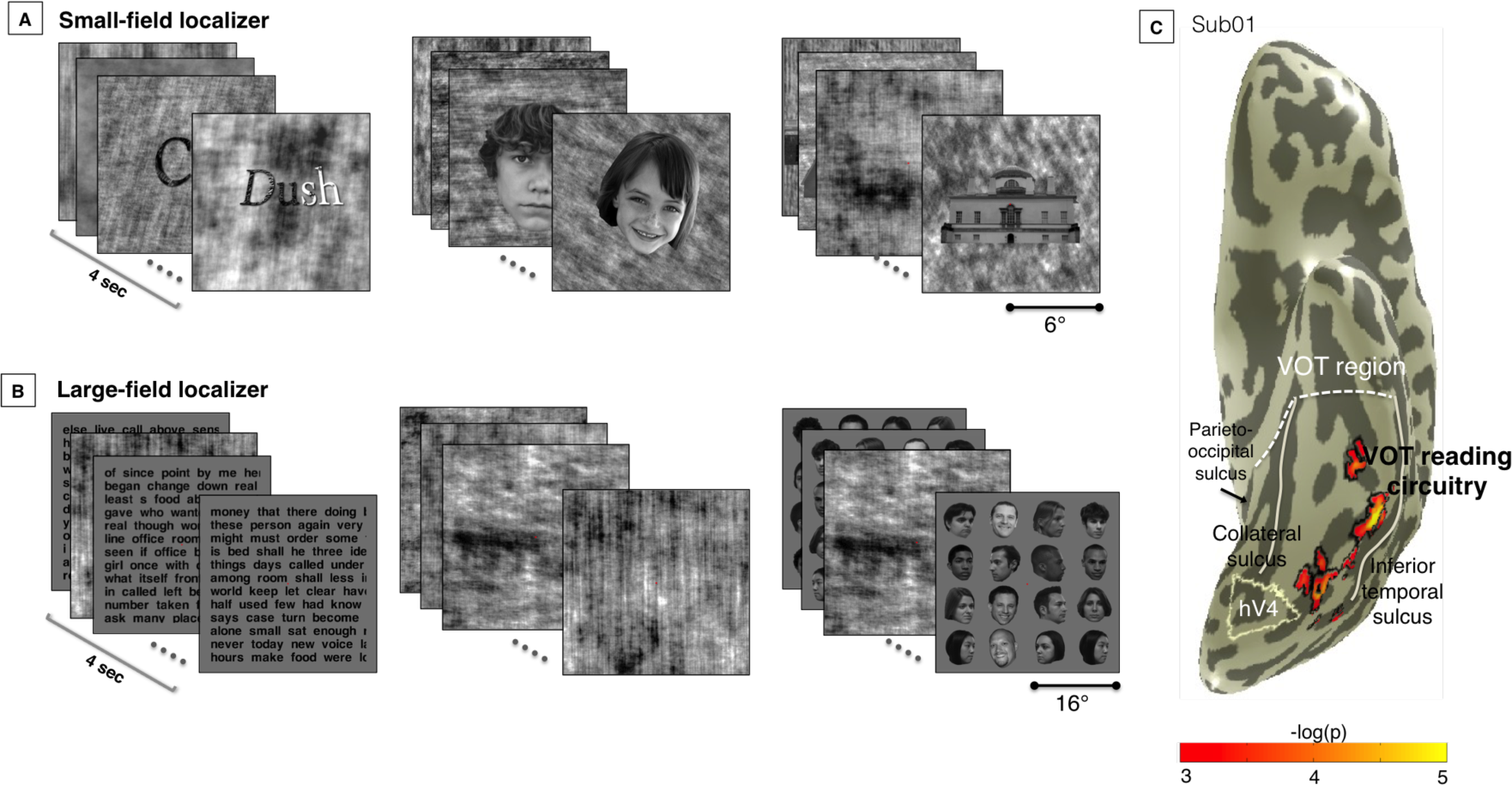
Functional and anatomical definition of the VOTRC. **A. The small-field localizer** presents a series of images from a single category, such as words, faces, objects and phase-scrambled objects. Each image is presented within the central 10° of the visual field. **B. The large-field localizer** presents a series of stimuli that each span the extent of the screen (32°). Stimuli are tiled with words or faces, or contain phase scrambled versions of the other categories. **C. The VOTRC** is the set of voxels (not-necessarily contiguous) within the VOT that are more responsive to words than to other categories. The VOT is bounded medially by the collateral sulcus, laterally by the inferior temporal sulcus, posteriorly by hV4, and anteriorly by an imaginary line drawn from the collateral sulcus to the inferior temporal sulcus, starting at the point where the parieto-occipital sulcus meets the collateral sulcus. Shown here is Sub01, large-field localizer.

#### PRF mapping

Stimuli for the the pRF mapping experiment were presented using the Eiki LC-WUL100L projector and controlled by code in the Psychophysics Toolbox (Brainard, 1997). A small dot (0.15° visual angle in diameter) at the center of the screen served as the fixation point. Participants were instructed to fixate the dot and to press a button when the dot changed color (randomly between red, green, and black every 1-5 seconds). Participants maintained fixation as a moving bar swept through the visual field in 4 orientations (0°, 45°, 90°, and 135° from vertical) and 2 motion directions. Eye-tracking was not used, but sharp hemifield and quarterfield pRFs in early visual field maps (Figure 2B, Figure 5CD) suggest that stable fixation was maintained by participants.

**Figure 5.**
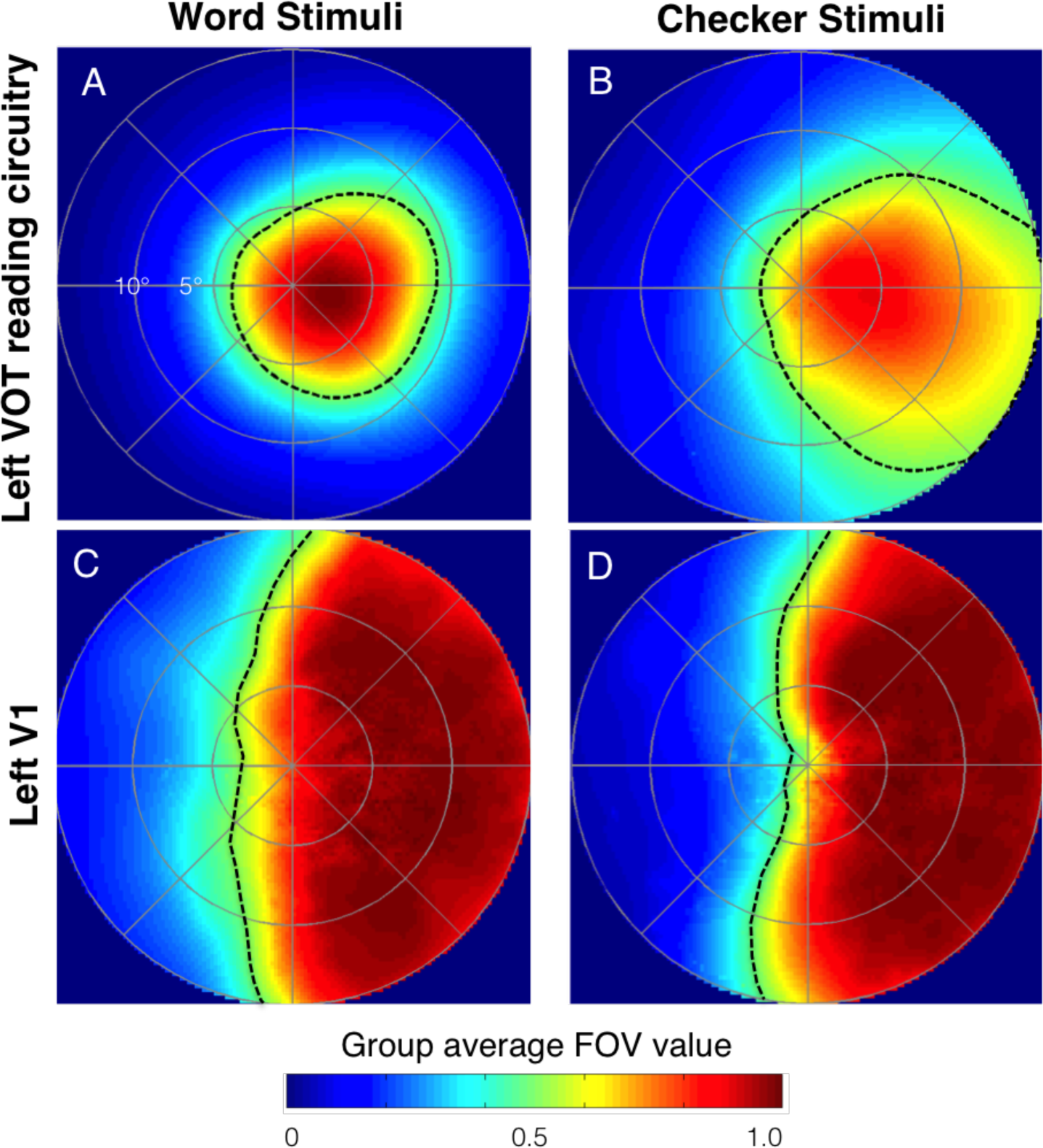
Stimulus dependence in word-responsive regions. Panels show the group average FOV (n=20) for the left VOTRC (A-B) and left V1 (C-D). The FOV in 5A is identical to the one shown in 3A. Population RF measurements made using word stimuli are shown on the left (A, C). Population RF measurements made using checkerboard stimuli are shown on the right (B, D). The dashed lines are the half-max contour of the group average. The FOV for each stimulus is estimated using only voxels where the pRF model explains at least 20% of the variance. For this reason, voxels included in the calculation of the FOV shown in A and B overlap, but are not identical. See Figure 6C for a comparison of group average FOVs within identical voxels.

In one set of runs, checkerboard stimuli were used as the contrast pattern within the bar. The checkerboards within the bars drifted parallel to the orientation of the bar and had a contrast reversal rate of 2 Hz. The size of each checkerboard square was approximately 1.3° per side. The checkerboard stimuli are described in more detail in previous work (Amano, Wandell, & Dumoulin, 2009; Dumoulin & Wandell, 2008). Each run was 192 seconds long. Three runs of checkerboard retinotopy were collected for each subject. Four blank periods (12s each) were interleaved to estimate larger pRF sizes (Dumoulin & Wandell, 2008).

In the other set of runs, word stimuli were used as the contrast pattern within the bar to better drive responses in the VOTRC. The words were rendered in black Helvetica font on a white background. The vertical extent of the lower-case letter ‘x’ was 1.3°. One-to-six letter words with frequency greater than 200 per million were used as the lexicon; words were randomly chosen from the lexicon to create a page of text. The text within the aperture was refreshed at 4 Hz. Each run lasted 5 min and two runs were collected. The runs of word and checkerboard stimuli were interleaved, and the order balanced across subjects.

**Figure 2.**
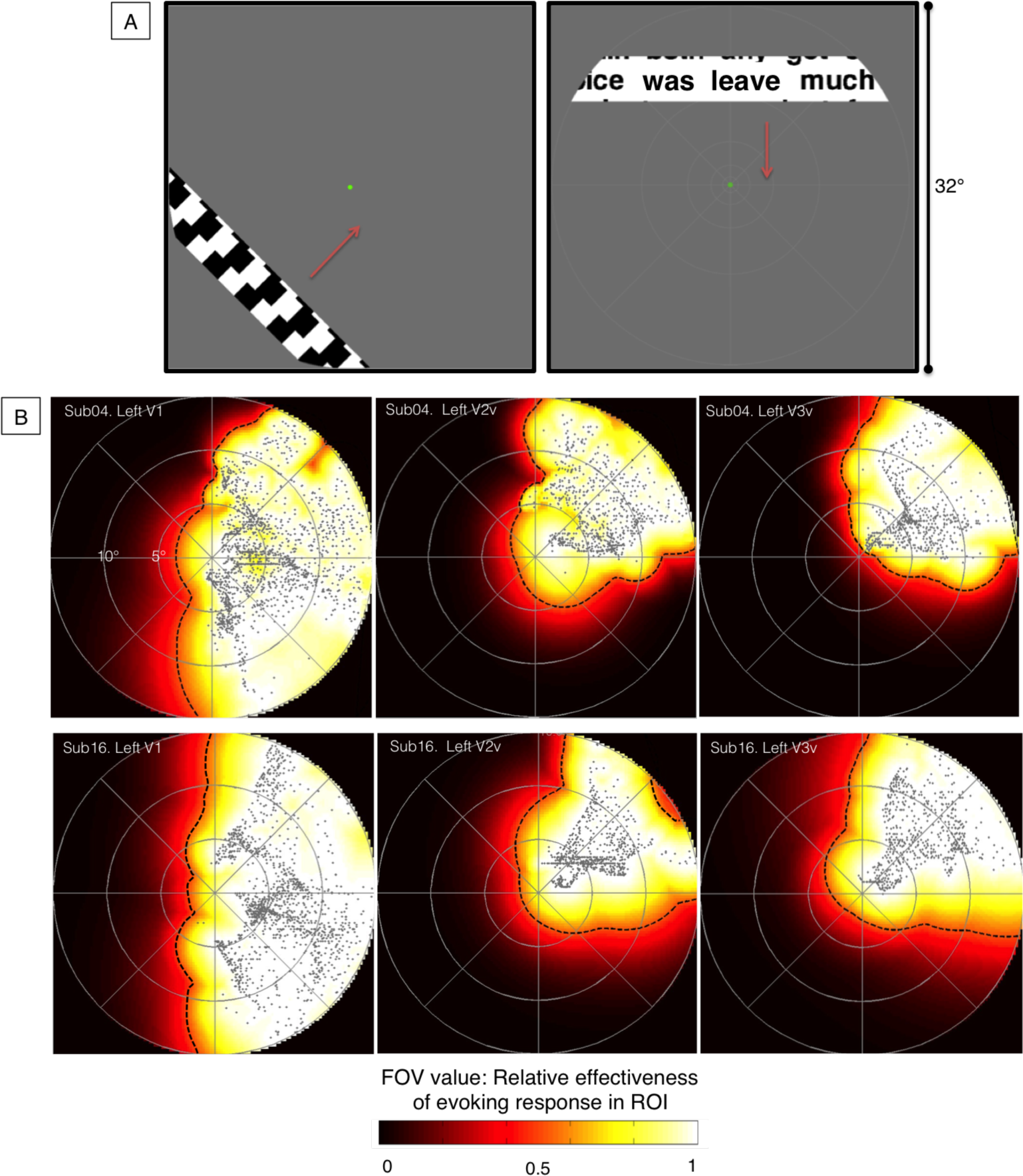
PRF mapping: Stimuli and visual field coverage. **A. Stimuli.** Stimuli consisted of bars (4° wide) that slowly and repeatedly traversed the visual field in eight different directions. The pattern within the bar contained dynamic checkerboards or words. **B. FOV for left V1, V2v and V3v from single subjects.** The points represent the pRF centers. The color indicates the largest pRF value across the collection of voxels in the ROI. This value can be interpreted as the relative effectiveness of stimuli evoking a response in the ROI. Population RFs were measured with word stimuli. Shown are the pRFs in V1 (left panels), V2v (middle panels) and V3v (right panels) of 2 participants, Sub04 (top row) and Sub16 (bottom row).

We model the population receptive field in each voxel using the compressive spatial summation (CSS) model (K. N. Kay et al., 2013). The CSS model estimates the pRF of a voxel by selecting parameters for a 2D Gaussian that optimally predicts the response to a translating stimulus (moving bars). The parameters of the pRF include location (x°,y°) and size (ρ°) in degrees of visual angle. The value of the 2D Gaussian at its peak is normalized to 1. The CSS model extends earlier linear models (Amano et al., 2009; Dumoulin & Wandell, 2008) by including a compressive nonlinear response exponent to account for subadditive responses which appear to increase across the visual hierarchy.

#### Estimating the FOV for a region of interest (ROI)

The FOV of a region of interest (ROI) defines the portion of the visual field that reliably evokes a response in any of the voxels in the ROI. The FOV of an ROI is obtained by aggregating over the voxels’ pRFs (the portion of the visual field where stimuli reliably drive responses). Each voxel has a pRF that is modeled as a 2D Gaussian in the visual field. The peak of the Gaussian is normalized to 1. The FOV of an ROI is obtained in the following way: at each point in the visual field, the FOV value is the maximum pRF value over the voxels. Unless otherwise stated, only voxels with greater than 20% variance explained by the pRF model are used in measuring the FOV.

To reduce the effect of pRFs that are anomalous in size and/or position, a bootstrapping procedure is applied (Winawer & Witthoft 2015, Winawer et al 2010, Amano et al 2009). Given a collection of N voxels in the subject’s ROI, N voxels are sampled with replacement, and the FOV is calculated for these N voxels. This procedure is repeated 50 times. The average of the 50 FOV samples is the FOV of the ROI. The FOV is a map of continuous values, so to characterize its size and shape, we describe the half-max contour, a contour encompassing the part of the visual field where the FOV values exceed 0.5. Example measurements showing the FOV and half-max contour are shown in Figure 2B (for V1, V2v, and V3v) and Figure 3B (for left VOTRC). The group summary FOV is obtained from averaging over the individuals’ FOVs (Figure 3C). In FOV measurements where voxels are thresholded at 20% variance explained, subjects with no voxels pass threshold are assigned a FOV of all zeros.

#### Quantifying the similarity between FOVs

The Dice similarity coefficient is used to quantify the similarity between FOVs. This is done by first representing the FOV as 128×128 matrix F, where each element of F can be thought of as a pixel with a value ranging between 0 and 1. A 128×128 binary matrix B is then obtained from F, where an element in B equals 1 if the corresponding pixel in F is greater than or equal to 0.5. In other words:

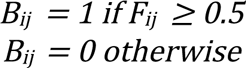

For binary matrices B1 and B2 from the two FOVs being compared, the Dice coefficient equals the number of pixel locations where both B1 and B2 equal 1 divided by the average of the number of pixels in B1 and B2 that equal 1. In other words:

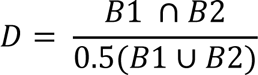

For two FOVs with exactly overlapping half-max contours, the Dice coefficient is 1. If the half-max contours do not overlap at all, the Dice coefficient is 0.

## Results

### The FOV of the VOTRC

We begin by considering FOV measurements made using words as the contrast pattern. The left VOTRC and the FOV for a single subject are shown in Figure 3A and B. The FOV is biased for the central portion of the contralateral visual field. In the typical subject, more than 98% of pRF centers (median across subjects, SD 11%) are located within 5 degrees of fixation. More than 97% of pRF centers (median across subjects, SD 18%) are located in the right visual field. The half-max contour extends further along the horizontal than the vertical meridian (Figure 3B). The FOV of the group (n=20) is shown in Figure 3C. The group average FOV demonstrates the same foveal and contralateral bias as the individual data.

**Figure 3.**
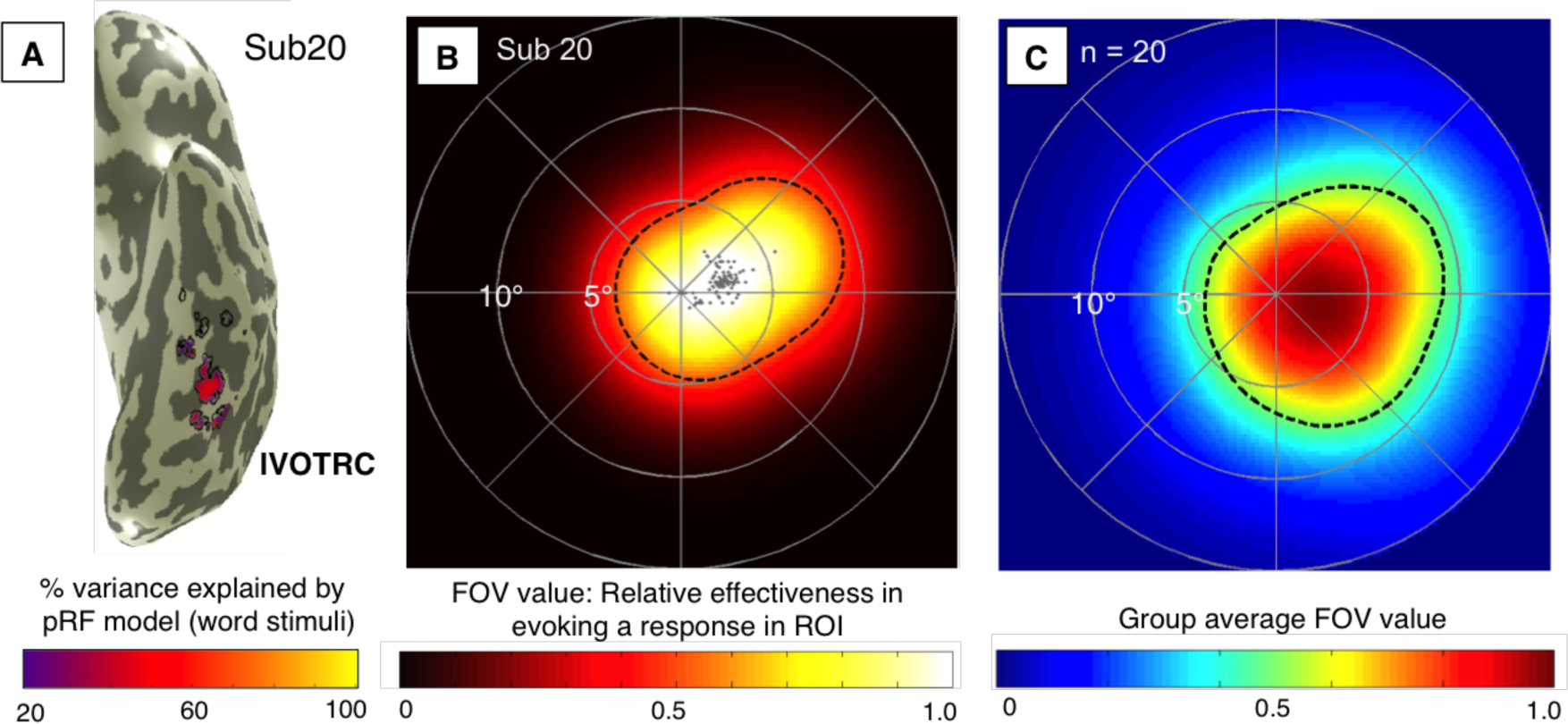
The FOV of the left VOT reading circuitry. **A. Left VOTRC definition.** A representative example of the left VOTRC (outlined in black) in a single subject. These voxels are more responsive to words than to other categories of visual stimuli (t-test, p < 0.001 uncorrected) in the large-field localizer (stimuli are shown in Figure 1B). Colors show the variance explained by the pRF fits using the CSS model with word stimuli. Colored voxels (with greater than 20% variance explained) were included in the FOV calculation. **B. The FOV of the left VOTRC.** Population RFs are estimated in response to word stimuli (the stimulus is shown in Figure 2A, right panel). The color map indicates the FOV value across the visual field. The dashed line is the half-max contour. **C. The group average FOV of the left VOTRC.** The average FOV of all 20 subjects is shown. The dashed line indicates the half-max contour of the average FOV.

### The FOV is reliable within subjects and varies between subjects

To assess within-subject reliability, we visualize the half-max contours from pRF models
fit to individual runs of data (Figure 4). For the most part, the independent half-max contours cover similar parts of the visual field (e.g., Sub04-07, Sub10-11, Sub16). In some cases the contours from the two runs differ substantially (e.g., Sub14, Sub20). In those cases, we typically find that one of the runs matches the FOV fit, and that this run was better fit by the pRF model. The data suggest that accurate FOV estimates require multiple (at least 2) independent runs.

The FOV of the left VOTRC varies between individuals. There are subjects in which both runs reveal small FOVs (Sub04) and others in which individual runs reveal large FOVs (Sub16). In other subjects the FOV seems reliably oriented in different directions (Sub05 and Sub06). Despite these differences, the half-max contour covers the ipsilateral foveal portion of the visual field for all subjects. The median between-subject Dice coefficient is 0.62 (minimum 0.26, maximum 0.88).

**Figure 4.**
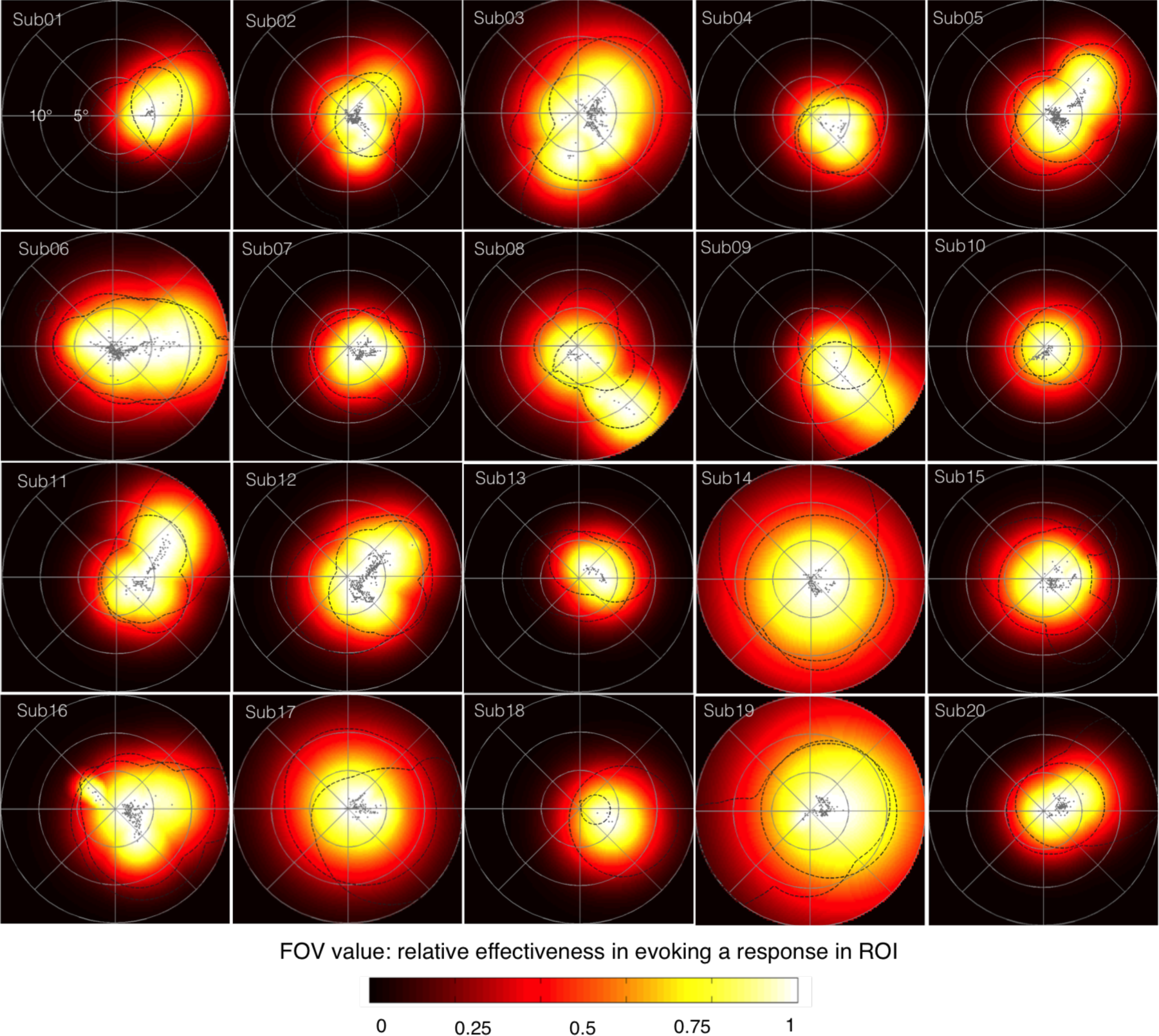
Field of view for the left VOTRC differs between subjects (n=20). Each panel depicts the FOV of the left VOTRC of a single participant. The color map and the gray dots (representing pRF centers) are calculated using voxels that exceed 20% variance explained from the pRF model fit to two runs of word stimuli. For the same voxels, the dashed lines outline the half-max contours from pRF models fit independently to run 1 and run 2.

### The FOV of the VOTRC, but not V1, is stimulus dependent

We measured pRFs using two different types of stimuli within the moving bar aperture: checkerboards and words. When measured with words, the average FOV of the left VOTRC extends 9° along the horizontal meridian in the contralateral visual field and 4° in the ipsilateral visual field (N=20, Figure 5A). When measured with checkerboard stimuli, the half-max contour of these same voxels extends further into the periphery but remains biased for the horizontal meridian (Figure 5B). The FOV does not change with the stimulus in early visual cortex. For example, the FOV of left V1 covers the contralateral hemifield when measured with words or checkerboard stimuli (Figure 5CD), in agreement with Winawer & Witthoft (2015) and Amano (2009).

### Stimulus dependency of the FOV is not explained by a difference in voxel selection or model accuracy

We consider two methodological explanations for the stimulus dependence of the FOV in left VOTRC (Figure 5A vs. 5B). One explanation may be due to the fact that non-identical voxels in left VOTRC pass the 20% variance explained threshold for each stimulus type. This means that different voxels are used in the calculation of the FOV in Figures 5A and 5B. However, choosing identical voxels for the calculation of the FOV results in the same stimulus dependence (see Figure 6C and Supplementary 4 and 5), thus the stimulus dependency is not due to the inclusion of different voxels for each stimulus type. Another explanation for the stimulus dependence may be due to differences in model accuracy: the pRF model explains the responses to words more accurately compared to checkerboards (Figure 6A). To investigate the possibility of the stimulus dependency being due to decreased model accuracy, we added Gaussian noise (~N(0,1.5)), to the left VOTRC responses (dashed blue lines, Figure 6B) and refit the pRF model. After adding noise, the mean model accuracy for this noisy time series is less than the mean model accuracy for the original word time series and the checkerboard time series (Figure 6A). We then estimated the FOVs for the noisy time series in the same set of voxels that were used to calculate the word FOVs of each participant. We find that the group averaged FOV estimated using the words+noise time course is very similar to the FOV obtained using the word responses (Dice coefficient 0.97, Figure 6C). This simulation demonstrates that the stimulus dependency on words and checkerboards is not explained by lower model accuracy and that the model is relatively robust to noise.

**Figure 6.**
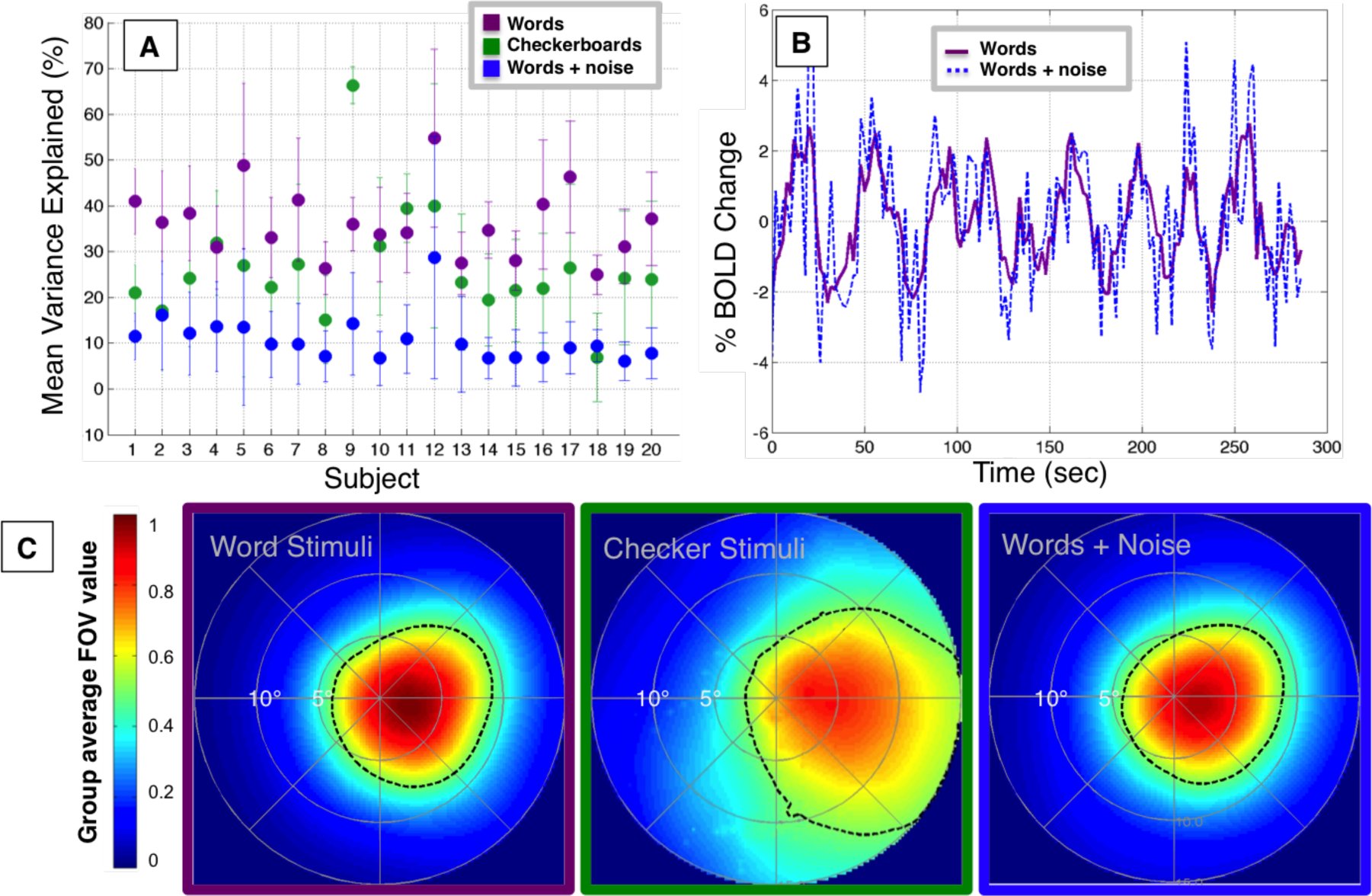
Noise simulation. **A. Variance explained.** Purple dots represents the mean variance explained in the left VOTRC for individual subjects, restricted to the voxels that have > 20% variance explained when measured with word stimuli. Error bars represent the standard deviation over voxels. Green dots show the variance explained in the same voxels when measured using checkerboards. Blue dots show the variance explained when Gaussian noise (s.d. 1.5%) is added to these voxels. **B. Example time series.** The time series of a voxel in the left VOTRC of Sub01 averaged over two runs as the subject viewed moving bars with words in the aperture (purple). The time series with artificially introduced Gaussian noise (s.d. 1.5%, dashed blue line). **C. Group averaged FOVs of left VOTRC.** Each panel shows the FOV for a different condition: words (left, same as Figure 3A), checkerboards (middle), and words+noise (right). The dashed contours indicate the half-max of the group average. These analyses are restricted to voxels that exceed 20% variance explained when measured with word stimuli, so that the voxels for each condition are identical (See Supplementary 4 and 5 for the complementary analysis using checkerboard responses for voxel selection). Other details as in Figures 3C and 5.

### The FOV of VOTRC is similar for two different localizers

In the small-field functional localizer, word images appear within the central 8° of the visual field (collected on Subjects 1-12). Restricting the stimuli to a small portion of the visual field may bias the voxel selection process towards voxels with a foveal preference. To address this concern, we use a large-field localizer in 9 additional subjects (Subjects 1, 13-20). The large-field localizer contains words that tile the central 32° and should identify word-responsive voxels in VOT that have positional preferences anywhere within this part of the visual field.

The VOTRC defined with the large-field localizer is similar to the definition with the small-field localizer (see Supplementary 1). One subject took part in both localizers and the difference is roughly what we find for repeated measurements (Figure 7A). The mean surface area of word-responsive cortex in VOT is 31 mm2 for the small-field and 39 mm2 for the large-field functional localizer, both with a standard error near 8 mm2 (surface areas for each subject are listed in Supplementary 2B). The group average FOV estimated with the two localizers is also similar (Dice coefficient = 0.91). In both cases, the FOV of the left VOTRC is biased for the horizontal meridian, extending ~9° in the contralateral visual field and ~4° in the ipsilateral visual field (Figure 7B).

**Figure 7.**
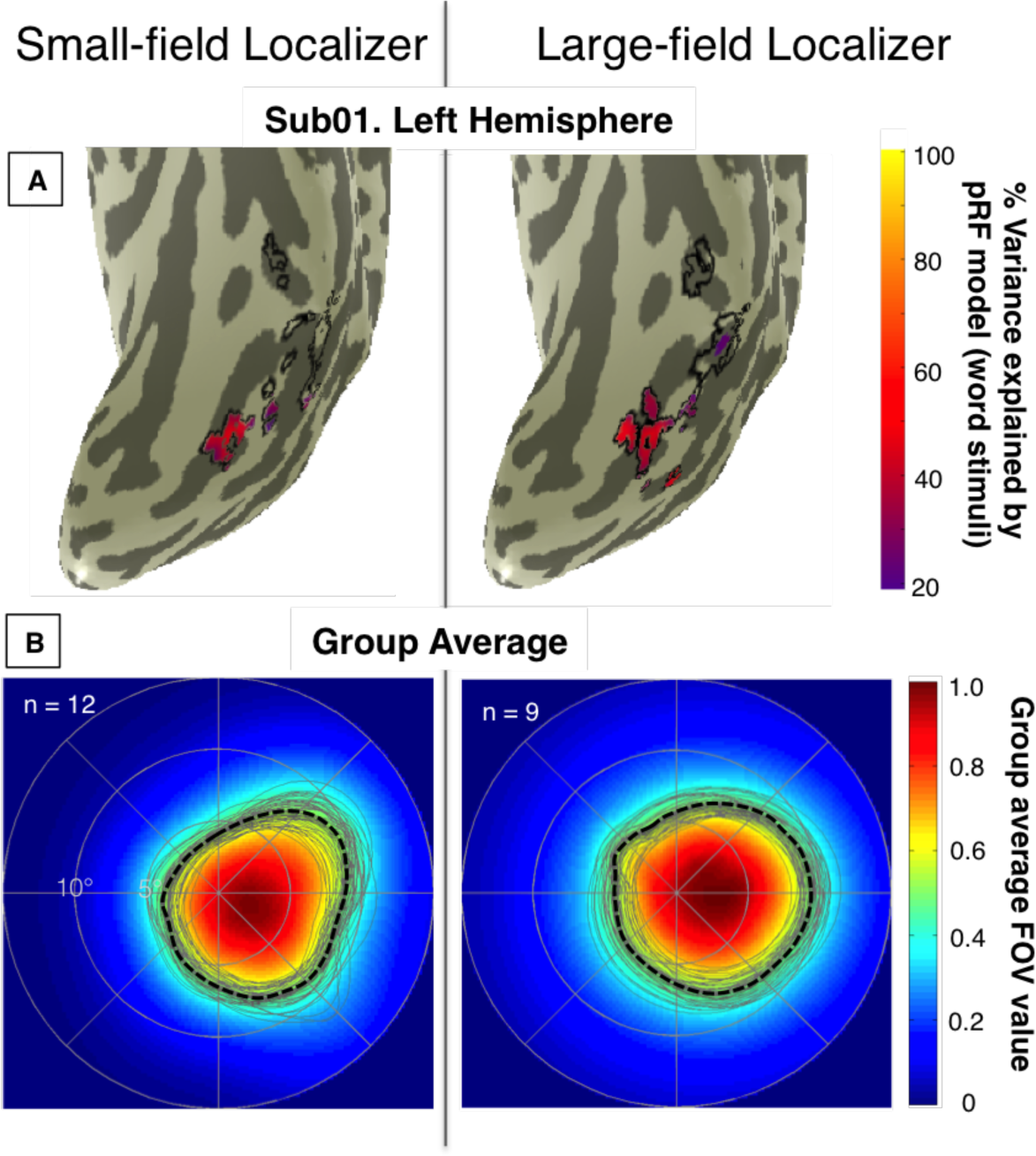
FOV comparison using the small- and large-field localizers. **A.** The left VOTRC (black outline) in Sub01 as defined by the 2 localizers: small-field (left) and large-field (right). Colored voxels have variance explained greater than 20% by the pRF model (word stimuli). **B.** The left panel shows the average FOV for Sub1-12 whose left VOTRC was defined using the small-field localizer. The right panel shows the average FOV for Sub1,13-20 whose left VOTRC was defined using the large-field localizer. The gray curves are 50 half-max contours of the bootstrapped group averages.

### The FOV of right VOTRC is contralaterally- and foveally- blased

The localizer experiments produced word-selective voxels in both hemispheres, allowing for the definition of a right VOTRC in every subject (M. Ben-Shachar, Dougherty, Deutsch, & Wandell, 2007; Cohen et al., 2002; Glezer, Jiang, & Riesenhuber, 2009; A. M. Rauschecker et al., 2012; Vigneau, Jobard, Mazoyer, & Tzourio-Mazoyer, 2005). Like the left VOTRC, the FOV of the right VOTRC exhibits a foveal and contralateral bias, extending 8° into the contralateral visual field and 3° into the ipsilateral visual field (Figure 8A). The Dice coefficient between the FOVs of the left and right VOTRC (when flipped) is 0.86; the two FOVs are mirror symmetric, with the right VOTRC FOV being slightly smaller (compare Figure 8B with Figure 5A, or see Supplementary 2A).

The distribution of pRF parameters in the left and right VOTRC are shown in Supplementary 2C-F. The principle difference is that there are more word-selective voxels in the left hemisphere (Supplementary 2B). Of the word-selective voxels, 49% in left VOTRC and 48% in right VOTRC have variance explained that exceeds 20%. The FOV of the combined left-right VOTRC covers an elliptical portion of the visual field that is centered on the fovea and extends mainly along the horizontal meridian (Figure 8D).

**Figure 8.**
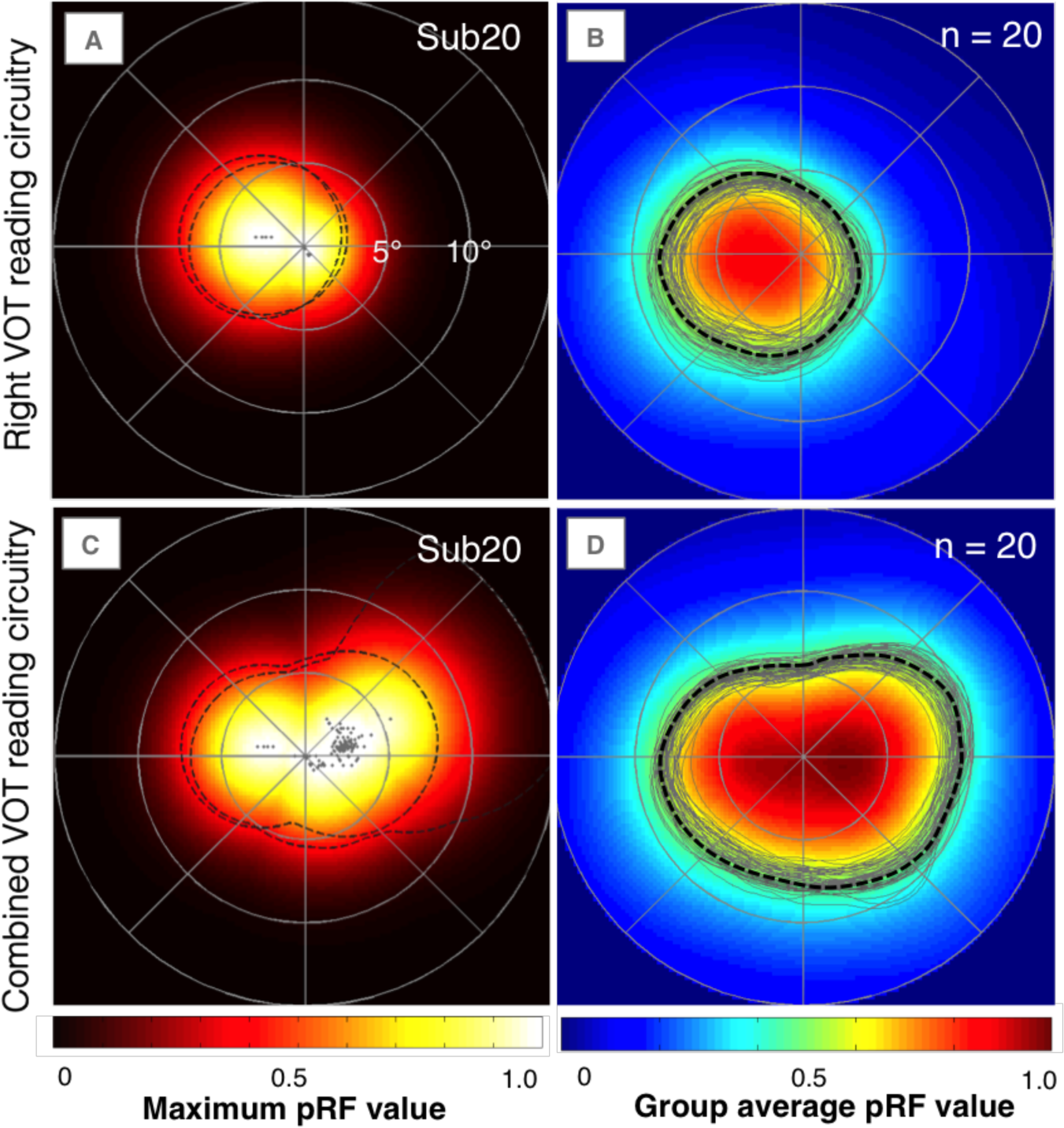
FOV of the right VOTRC and of the combined VOTRC. **A.** Right VOTRC FOV for a representative subject. **B.** Group average FOV of the right VOTRC. In one subject, no voxels passed threshold. **C.** Left and right (combined) VOTRC FOV for a representative subject. **D.** Group average FOV of the combined VOTRC. The dashed lines indicate the half-max contours of individual runs. Gray dots in A and C represent pRF centers in the model fit to the average of the two runs. The gray contours are the half-max of 50 bootstrapped group averages.

### The VOTRC’s FOV is restricted to a central region of the VOT’s FOV

While there is variation in the size and shape of the FOV between subjects, the FOV of the combined left and right VOTRC of the average subject has a distinctive shape. The average FOV extends about 9° along the horizontal meridian and about 6° along the vertical meridian. The FOVs of the left and right VOTRC are roughly mirror symmetric, although the FOV of the right VOTRC is slightly smaller. The FOV of the combined VOTRC is a fraction of the FOV of the full VOT, which extends farther into the periphery in both the horizontal and vertical directions (Figure 9).

**Figure 9.**
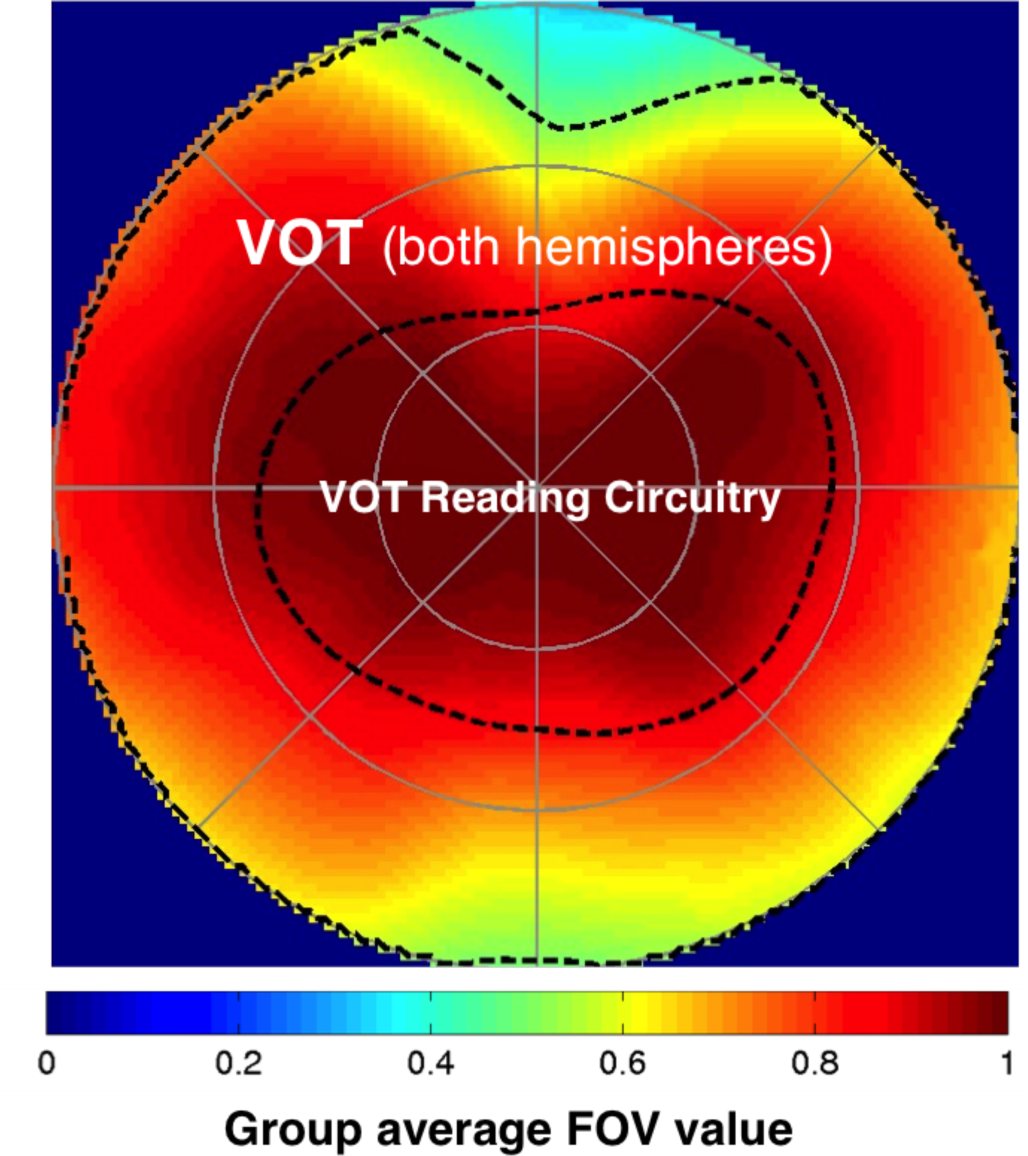
Comparison of FOVs for the VOT and the VOTRC. The dashed lines indicate half-max contours of the group average FOV. The outer contour and the color map show the FOV of the VOT, both hemispheres (Figure 1C). The inner contour shows the FOV of the combined VOTRC (Figure 8D).

### PRF model accuracy

Model accuracy is derived from two calculations that use the root mean squared error (RMSE)

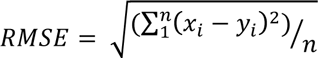

where n = 144 (number of time points) and xi and yi correspond to the values of the time series being compared at a specific time point i. The first RMSE calculation is between independent time series data collected in runs 1 and 2. We call this Datarmse, and it can be interpreted as a measure of test-retest reliability. The second RMSE calculation is between the the model prediction obtained from run 1 and the actual time series measured in run 2. We call this Modelrmse. Since Run 2 is independent of run 1, Modelrmse is a cross-validated error measure of the model prediction. Finally, Relativermse is the ratio of the two RMSE calculations: Modelrmse / Datarmsee and we use this ratio to summarize model accuracy (see Figure 9). All time series are responses obtained from pRF mapping with word stimuli.

More than 99% of the combined VOTRC voxels have a Relativermse < 1. This means the model predicts run 2 with higher accuracy (smaller error) than the test-retest reliability (Modelrmse is smaller than Datarmse). A model is better than test-retest reliability when it captures an essential part of the response and averages out measurement noise. If a model is a perfect description of the mean signal and the measurement noise at each time point is a Gaussian distribution, the Relativermse values should be at 1/sqrt(2) = 0.707 (Rokem et al., 2015). The median value of the Relativermse across all VOTRC voxels is 0.73, close to the level of a model that extracts all of the reliable information available in the data.

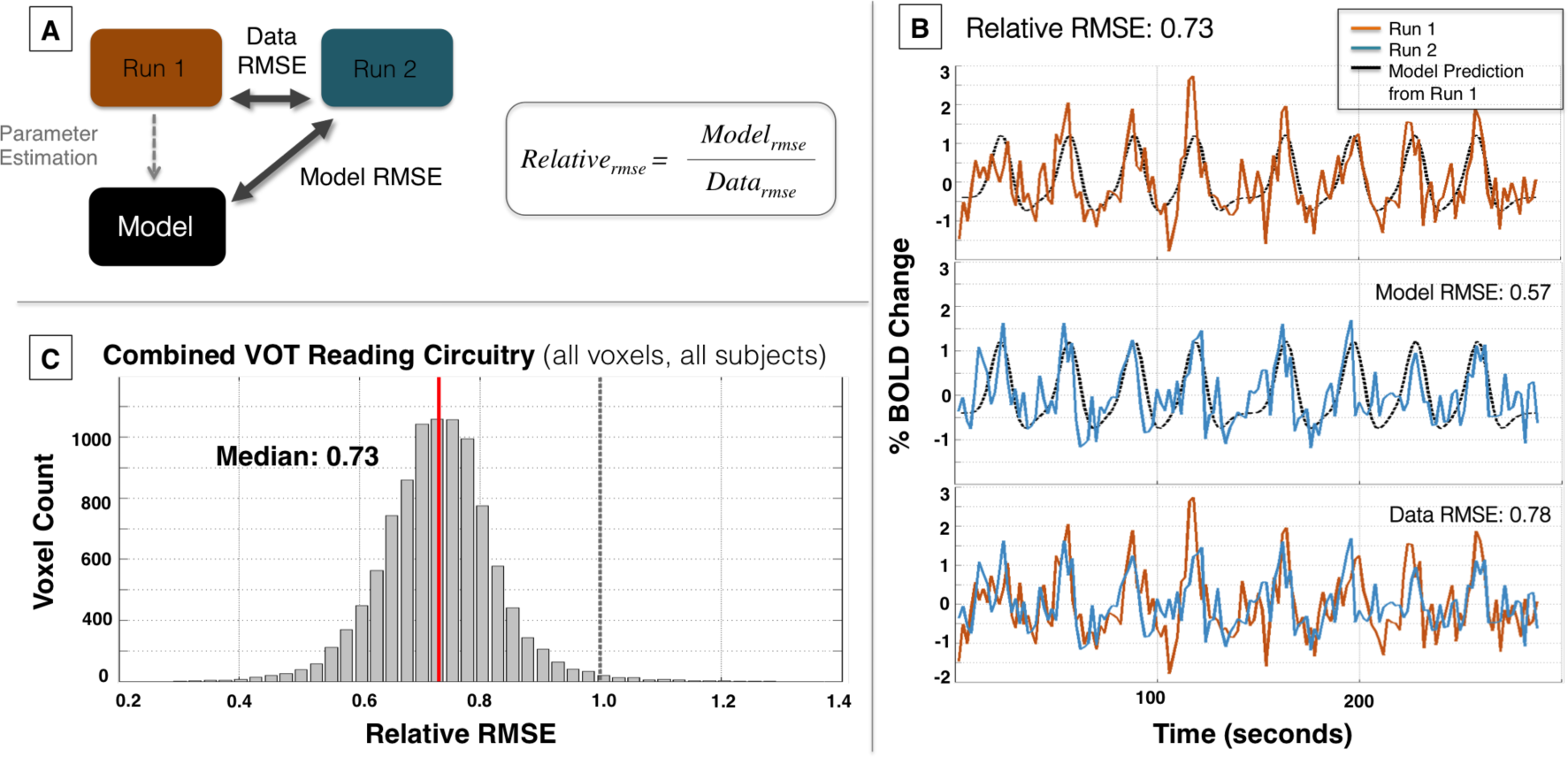

## Discussion

### Position sensitivity in the VOT

The VOTRC is embedded within a larger cortical territory (VOT) that contains several other functionally specialized regions (e.g. Epstein & Kanwisher, 1998; Grill-Spector & Weiner, 2014). We consider position sensitivity and FOV estimates of the VOT generally and then discuss specific features of the reading circuitry.

There is consensus that the functional responses of many VOT voxels are sensitive to stimulus visual field position (e.g., (Arcaro, McMains, Singer, & Kastner, 2009; Kravitz, Vinson, & Baker, 2008b; Uyar, Shomstein, Greenberg, & Behrmann, 2016). For example, more lateral VOT regions including the fusiform gyrus are more responsive to stimuli within 1.5° of the fovea while medial regions, including the collateral sulcus and lingual gyrus, are relatively more responsive to stimuli presented at greater than 7.5° eccentricity (see Figure 4 in Hasson, Levy, Behrmann, Hendler, & Malach, 2002. The VOT in general is more responsive to stimuli presented in the contralateral than ipsilateral visual field (Kravitz et al., 2008a; A. M. Rauschecker et al., 2012).

Several groups have used pRF mapping and FOV measurements to characterize extrastriate visual cortex (Amano et al., 2009; Larsson & Heeger, 2006; Brian A. Wandell & Winawer, 2011). A number of recent papers have specifically measured pRFs and FOV in VOT regions located near the reading circuitry (Kendrick N. Kay, Weiner, & Grill-Spector, 2015; Silson, Chan, Reynolds, Kravitz, & Baker, 2015). Silson et al. (2015) report that pRF centers in the occipital face area (OFA) and fusiform face area (FFA) are located within the central 5° and have a contralateral bias. They report that the FOVs in these two regions represent the lower (OFA) and upper (FFA) quarterfields. Kay et al. (2015) and Witthoft et al. (2016) agree that face-selective regions have a contralateral bias and that pRF centers are consistently near the fovea. They do not report the lower/upper field bias in apparently corresponding face-selective regions (IOG and FFA respectively).

The pRF model implementations and the cortical regions of interest differ between these three studies. Kay et al. use a model that includes a compressive nonlinearity (K. N. Kay et al., 2013). Silson et al. use a linear model implemented within AFNI, and Witthoft et al. use a linear model implemented in Vistasoft. In addition to the model differences, the anatomical definitions differ. Kay et al. and Witthoft et al. subdivide the FFA into two distinct regions, mid-fusiform (mFus) and posterior fusiform (pFus) (Weiner & Grill-Spector, 2012). Silson et al. (2015) refer to the OFA which may not be precisely the same as the IOG. Hence, the differences in findings, primarily the upper/lower visual field bias, may be due to the variation of methods used between groups. The contralateral and foveal biases that remain despite the methodological differences suggest that these are robust properties.

### Lateralization of the VOTRC

Most work on word-responsive regions in visual cortex focuses on the left hemisphere (e.g. Bouhali et al., 2014) because the responses in the left are more selective and larger than those in the right (Cohen et al., 2002; others). Additionally, there is a substantial neurological literature that associates alexia with lesions to the left hemisphere and emphasizes differences in representational capacity between the two hemispheres (Damasio & Damasio, 1983; Dehaene & Cohen, 2011; Dejerine, 1892). Nonetheless, it is important to recognize that word-selective responses are reliably observed in both hemispheres (M. Ben-Shachar et al., 2007; Cohen et al., 2002; Glezer et al., 2009; A. M. Rauschecker et al., 2012; Vigneau et al., 2005). Evidence in callosal patients suggests that the right hemisphere alone is sufficient for some reading functionality (Baynes, Tramo, & Gazzaniga, 1992). The left lateralization of responses emerges during development, and some authors propose that learning to read reduces word-selective responses in the right hemisphere (Shaywitz et al., 2007; Simos, Breier, Fletcher, Bergman, & Papanicolaou, 2000). The FOV of the left VOTRC is approximately mirror symmetric to the right VOTRC (Figure 8CD). The principal difference is that there are more word-selective voxels in the left hemisphere (Supplementary 2B).

Some earlier models of object recognition (Riesenhuber & Poggio, 1999; Rolls, 2000) and earlier experiments involving the visual word form area (VWFA; (Cohen et al., 2002; Dehaene & Cohen, 2011) report that left hemisphere responses to words are invariant to position in the visual field. This summary is at odds with the contralateral bias of the FOV described here. But examination of the data in these earlier experiments shows that responses in the VWFA are indeed elevated for words presented in the right visual field (e.g., Figure 5 in Cohen et al., 2002, also see Figure 5 in M. Ben-Shachar et al., 2007; Figure 5 in (A. M. Rauschecker et al., 2012). Hence, the findings here are consistent with those reports, and we extend them by establishing a quantitative FOV measurement.

### Stimulus dependency within the VOT

Using checkerboards rather than words within the aperture decreases the variance explained by the pRF model and increases the FOV (Figure 5). This effect is not well explained by methodological considerations, as we showed in detail (Figure 6, Supplementary 4 and 5). We discuss two possible explanations of the stimulus dependence of the FOV.

One possibility is that word stimuli and checkerboard stimuli are carried to the VOT through different pathways that carry information from slightly different FOVs. For example, Rauschecker et al. (2011) proposed that pattern contrast and motion contrast are communicated to the VOTRC via different routes. An alternative possibility is that different neuronal populations within the voxel preferentially respond to checkerboard and word stimuli (Harvey, Fracasso, Petridou, & Dumoulin, 2015). Stimulus dependence could occur if these populations have different FOVs.

Currently our model represents the input stimulus as a binary mask that indicates where there is some stimulus contrast (i.e. only indicating the position of the moving bar and not the image within the bar). A model that takes into account the image within the moving bar may be able to account for some of the stimulus dependency. Further, it may be possible to create circuit models that account for the contributions of multiple cortical processing regions (K. Kay & Yeatman, 2016); such models may also account for the stimulus dependence.

### Relationship to psychophysical measures of reading: Visual and perceptual span

The data suggest that the relatively large differences we observe between the FOVs in different subjects are reliable (Figure 4, compare Sub13 and Sub16). The FOV measurements describe a characteristic of the reading circuitry, and the size and shape of the FOV may be related to certain psychophysical measures of reading. We suspect there may be a compliance range for the field of view in the VOTRC, such that an inadequate (too small) or an irrelevant (too large) FOV disrupts processes that are necessary for learning to rapidly recognize words. If this is true, then the FOV will be predictive of some reading-related measures. Two candidate psychophysical measures that may be predicted by the shape and size of the FOV are the visual span (Gordon E. Legge, Ahn, Klitz, & Luebker, 1997) and the perceptual span (McConkie & Rayner, 1975).

The visual span is the number of letters of a given size, arranged horizontally, that can be reliably recognized without moving the eyes. In typical readers of English under typical reading conditions, the expected range of visual span values is about 7-11 letters (Gordon E. Legge et al., 1997). Individual differences in reading speed depend on multiple factors, and the visual span positively correlates with reading speed and accounts for a 34-52% of this variation (Kwon, Legge, & Dubbels, 2007; G. E. Legge et al., 2007). The visual span can also be thought of as the number of letters that are not crowded (Pelli et al., 2007). At the age where children learn to read, the crowding effect decreases with age (Bondarko & Semenov, 2004).

The perceptual span, assesses reading performance while allowing for the influence of contextual information and linguistic factors. The perceptual span is defined as the region of the visual field available to the reader at a single fixation which allows the individual to maintain their typical reading speed (McConkie & Rayner, 1975). In typical English readers, the perceptual span encompasses about 15 characters to the right of fixation and 3-4 characters to the left of fixation (McConkie & Rayner, 1975). O’Regan (1991) describes the difference between perceptual span and visual span: “…‘visual span’ refers to what can be seen without the help of linguistic knowledge or context, whereas perceptual span includes what can be seen with that help”. The visual and perceptual spans are often studied separately but it seems likely that the size of the perceptual span is influenced by the size of the visual span.

Variations of the input from early visual cortex to the reading circuitry may influence reading performance. The between-subject differences in the FOV (Figure 4) may predict individual differences in visual span or even reading speed at different locations in the visual field. However, linking the FOV to behavior will require more theoretical development and additional measurements. As an example of the complexity, note that other systems (such as eye movements, language and memory) contribute to reading so that differences in the FOV may not map directly to differences in reading performance.

### Implications for reading development for impaired and non-impaired readers

The VOTRC represents a restricted subset of the information available from earlier retinotopic cortical regions (Brian A. Wandell et al., 2012). From a developmental perspective, one possibility is that the restricted FOV is present even prior to print exposure. Another possibility is that the foveal and horizontal bias in the FOV of the VOTRC is carved out during the early school years, as the VOT circuitry develops enhanced sensitivity to written words (Michal Ben-Shachar, Dougherty, Deutsch, & Wandell, 2011b). These two hypotheses could be discriminated by measuring the FOV in pre-reading children (4-5y) using words and checkerboards. If the FOV develops through education, the word FOV in pre-readers will resemble the checkerboard FOV in both groups. With greater exposure to print, the word-FOV may show increased bias for the central and horizontal parts of the visual field.

The reduction of the word-FOV may reduce irrelevant signals to the reading circuitry and improve word-processing efficiency (Kwon et al., 2007). On the other hand, a FOV that is too small may constrain the use of direct access, holistic strategies for visual word recognition (Coltheart, Rastle, Perry, Langdon, & Ziegler, 2001). This may lead to slower, piecemeal recognition of written words, particularly for longer words. Indeed, some dyslexics exhibit letter-by-letter reading (Wimmer & Schurz, 2010), and it may be the case that such behavior is correlated with a reduced FOV in the VOTRC. Importantly, reading acquisition is clearly limited by other factors, most prominently phonological processing (e.g., Ziegler & Goswami, 2005). Reading relies on multiple brain circuits that may be constrained or impaired in multiple ways, giving rise to the highly variable individual profiles of poor readers (Michal Ben-Shachar, Dougherty, & Wandell, 2007; Stanovich, 1986; Brian A. Wandell & Yeatman, 2013).

### Limitations

These measurements and analyses have both biological and experimental limitations. The BOLD measurements are limited by the presence of large sinuses in the neighborhood of the ventral occipito-temporal region. The transverse sinus interferes with the functional responses, which may have an impact on the FOV measurements (Winawer, Horiguchi, Sayres, Amano, & Wandell, 2010). The variable position of this sinus across participants may account for some of the between-subject variation that we observe in the FOV of the reading circuitry.

A very large number of experimental parameter choices must be made in designing the localizer. In some cases, the choice does not alter the FOV (small-field vs. large-field localizer). In other cases, the FOV measurements may be less robust to experimental changes. Experimental parameters include stimulus categories, baseline stimuli, and task requirements. Additional experimental choices are made in the design of the pRF portion of the experiment. Here, relevant parameters include the size of the words, the size of the bar aperture, and the rate at which the words refresh within the aperture, in addition to the task requirements. Further experiments will be needed to assess the effect of these choices on the FOV estimates.

Data analysis introduces further choices. In defining the VOTRC, we select voxels exceeding an uncorrected statistical threshold of p<0.001. In calculating the FOV in response to word stimuli, we include voxels for which the pRF model explains greater than 20% of the variance in the data. These parameters select the voxels with the largest and best-modeled responses, but there are many voxels with smaller responses that may contribute to reading performance. The results we report are robust to a wide range of parameters at the level of the group average, but individual FOV estimates may be sensitive to these parameters in a way that is yet to be determined in future studies.

## Conclusion

The FOV of the VOTRC is a small subset of the entire field of view available to the human visual system. On average, the FOV extends about 9° into the contralateral visual field and 4° into the ipsilateral visual field, and is largely confined to the horizontal meridian in each visual field. Unlike V1, FOV measurements in the VOTRC depend on stimulus features, and the FOV is more foveally biased for words than checkerboards.

In VOT broadly, the size and center of pRFs depend on factors such as attention, task requirements, and stimulus structure (e.g. the size of the words in the aperture) (Harvey et al., 2015; Kendrick N. Kay et al., 2015; K. N. Kay et al., 2013; Silson et al., 2015; Sprague & Serences, 2013). The restricted FOV of the VOT circuitry may be determined by the properties of the neurons within the VOT or the major projections to these neurons from other parts of cortex. It is also possible that voxels contain multiple populations of neurons with different sensitivities (Harvey et al., 2015). Current pRF models do not capture such stimulus and task sensitivities, and thus a next generation of models may be required to explain the broader properties of the VOTRC responses (K. Kay & Yeatman, 2016). We have archived the experimental data and metadata in a sharable database for anyone who would like to carry out further analyses (B. A. Wandell, Rokem, Perry, Schaefer, & Dougherty, 2015).

The properties of the neurons and projections in VOT may develop in response to the extensive training that many children undergo in school. This training may impact several systems, including the VOT reading circuitry, eye-movement circuitry, and connections to
language cortex. There may be multiple system solutions for normal reading performance; a limitation in one part of the circuitry (e.g. a small FOV) may be compensated by another part of the reading circuitry (e.g. a more efficient eye-movement system). The timing of the training with respect to the available plasticity in these different systems may matter for these learning processes. We hope that understanding the developmental processes in the reading circuitry will contribute to effective educational practice (Yeatman, Dougherty, Ben-Shachar, & Wandell, 2012).

## Acknowledgements

This work was supported by NSF Grant to BW and the BSF Grant to MBS and BW. We thank Michael Barnett for his help with data collection as well as construction of the display in the magnet bore. We thank Justin Gardner, Karen Larocque, Kendrick Kay, Jason Yeatman, Jon Winawer, and Lee M. Perry for their helpful comments and feedback.

## Appendix

**Supplementary 1.**
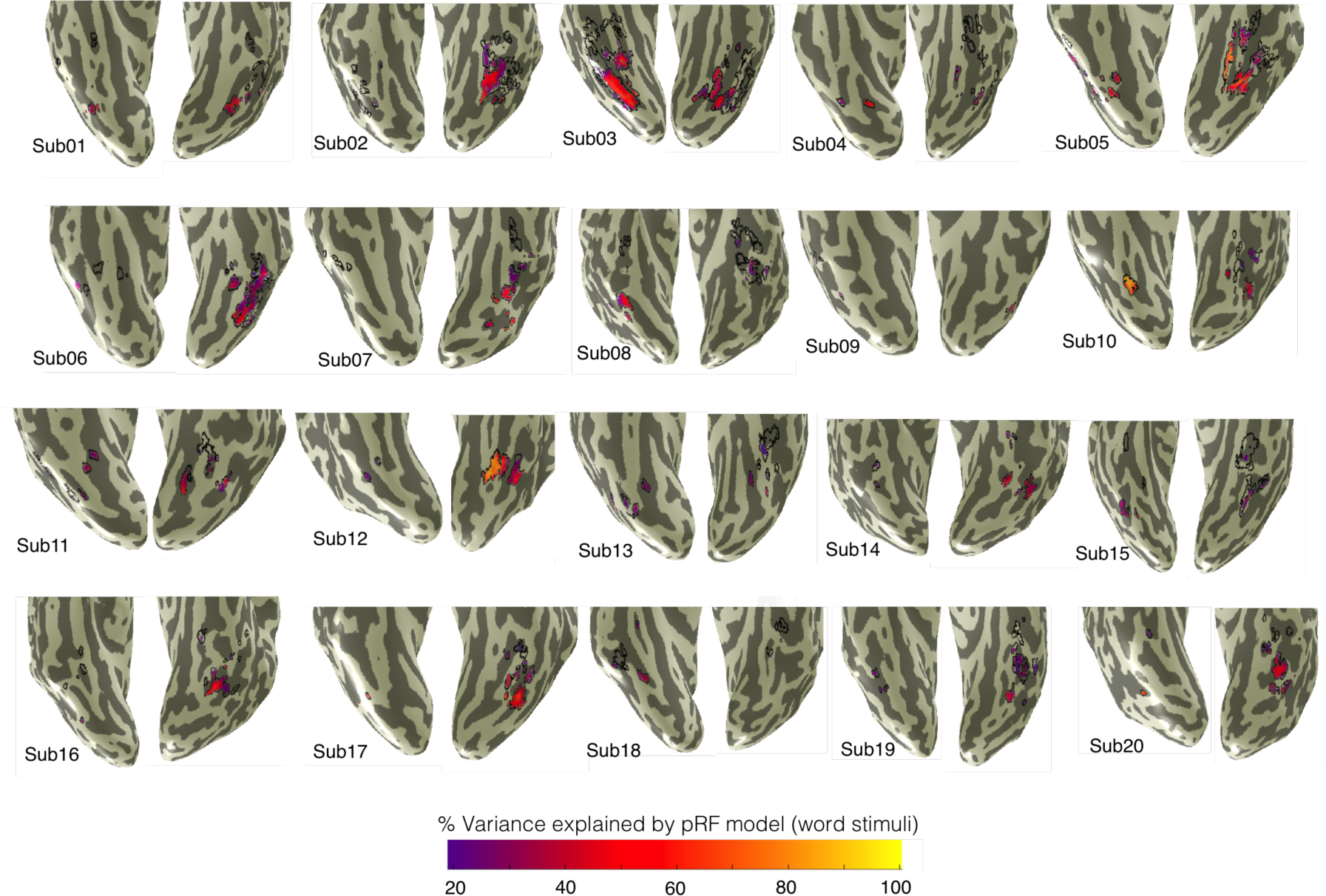
VOTRC definition for all subjects. The VOTRC is defined as those voxels within an anatomical region that are more responsive to words than to other categories. The black outlines show the left and right VOTRC in 20 subjects. In Sub 1-12 the regions were obtained with the small-field localizer; in Sub 13-20 the regions were measured with the large field localizer. The color overlays show the variance explained by the pRF model when using word stimuli. Only voxels with responses that exceed 20% variance explained are colored.

**Supplementary 2.**
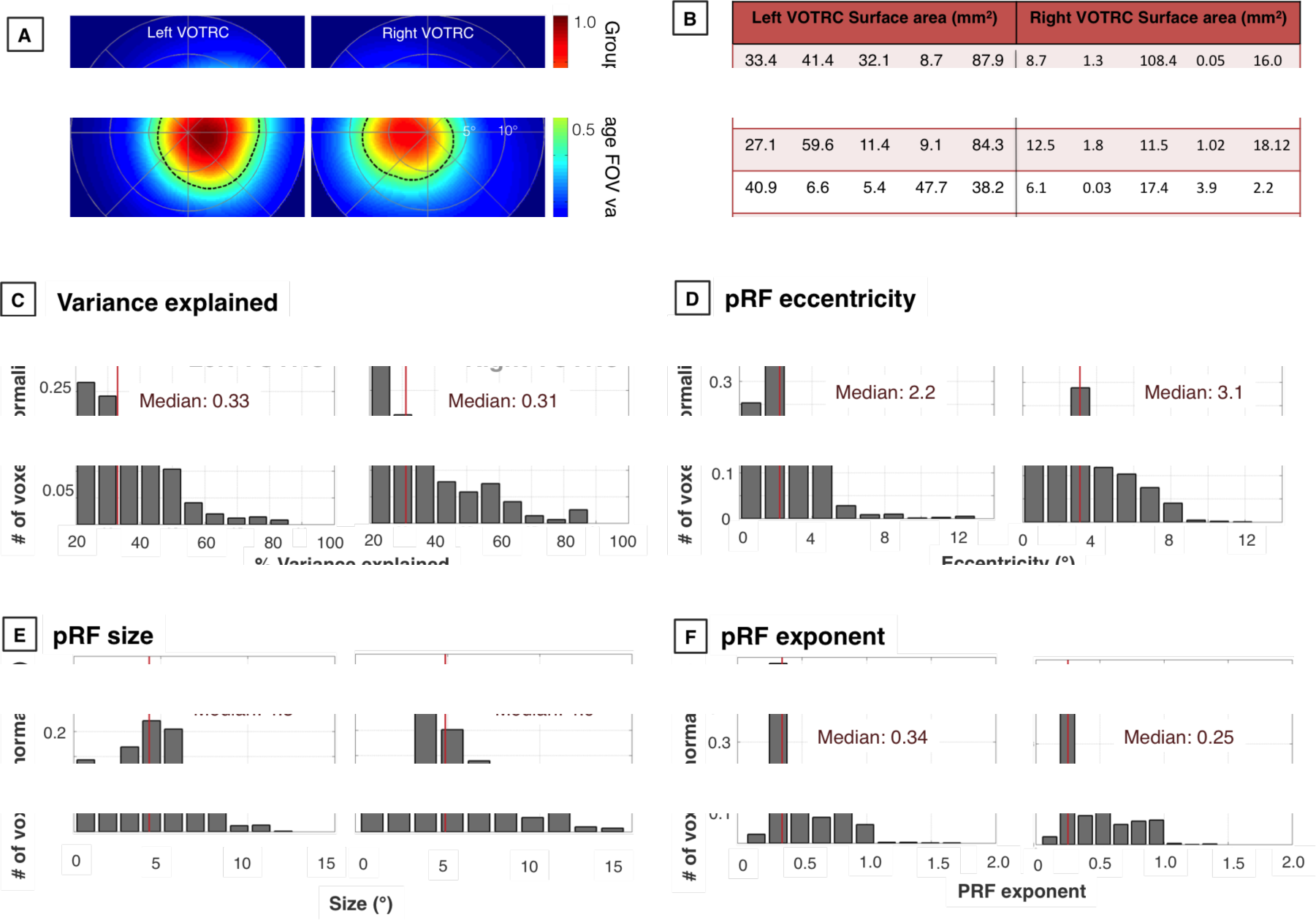
Distribution of pRF parameters. **A.** Group average (n=20) FOV of left and right VOTRC. Dashed curves are the half-max contour. **B.** Surface area (mm^2^) of the left and right VOTRC for individual subjects. Subject numbering follows the convention of Figure 4. **C,D,E, F.** Distributions of pRF parameters. 200 voxels are sampled with replacement from each subject and plotted for left and right VOTRC. Only voxels that exceed 20% variance explained are included.

**Supplementary 3.**
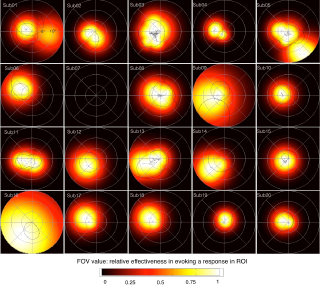
(Related to Figure 4). Between- and within-subject reliability in the right VOTRC. Dashed lines indicate the half-max contour to pRF models fit to run 1 and run 2. The color map and the centers correspond to the pRF model that is fit to the average of the two runs. For subject 07 no right hemisphere voxels passed the threshold criterion.

**Supplementary 4.**
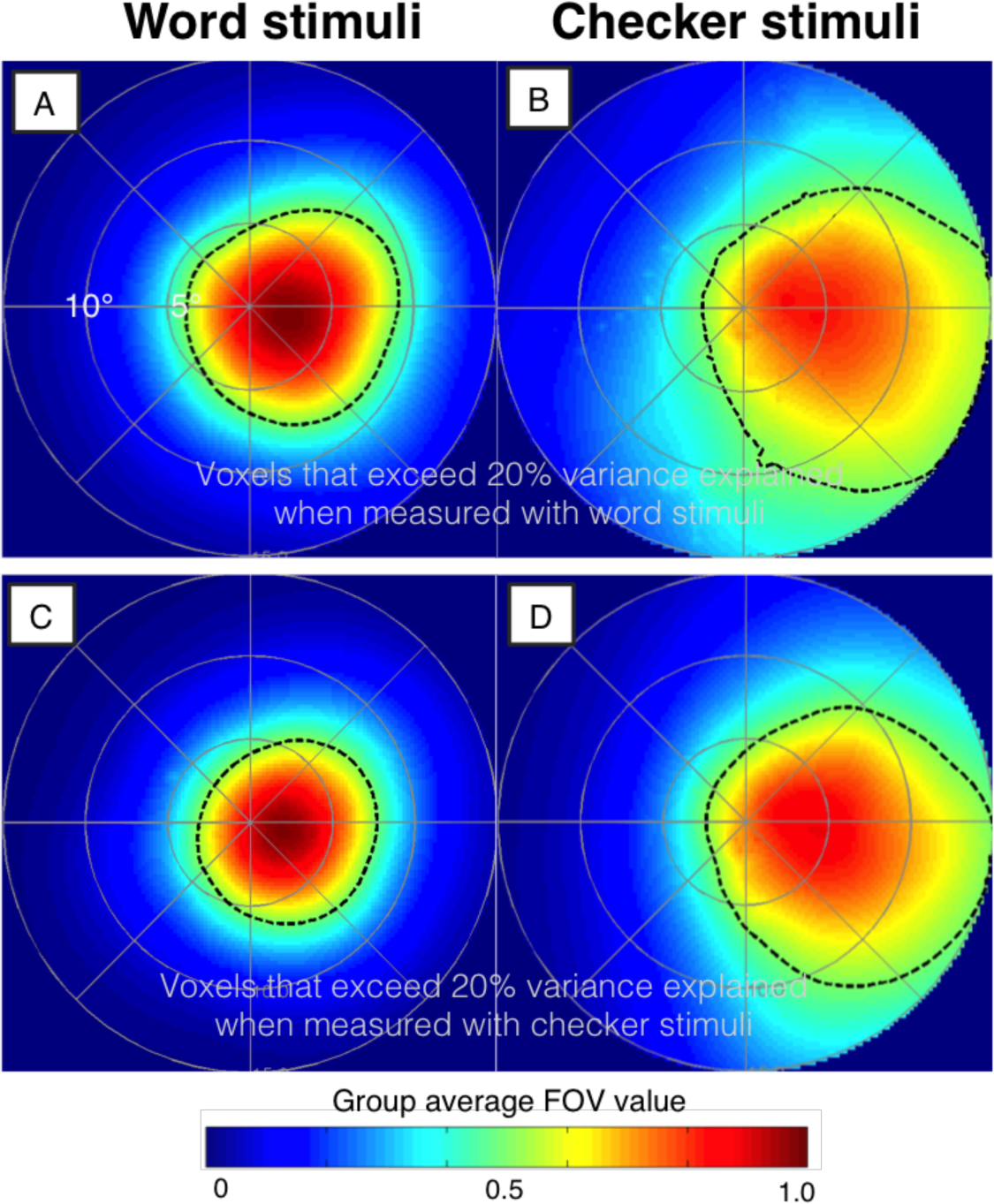
Group average FOVs in left VOTRC for various subset of voxels. FOVs in A and C were measured in response to word stimuli. FOVs in B and D were measured in response to checker stimuli. The left VOTRC voxels used in A and B are those that exceed 20% variance explained when measured with word stimuli. The left VOTRC voxels in C and D are those that exceed 20% variance explained when measured with checker stimuli.

**Supplementary 5.**
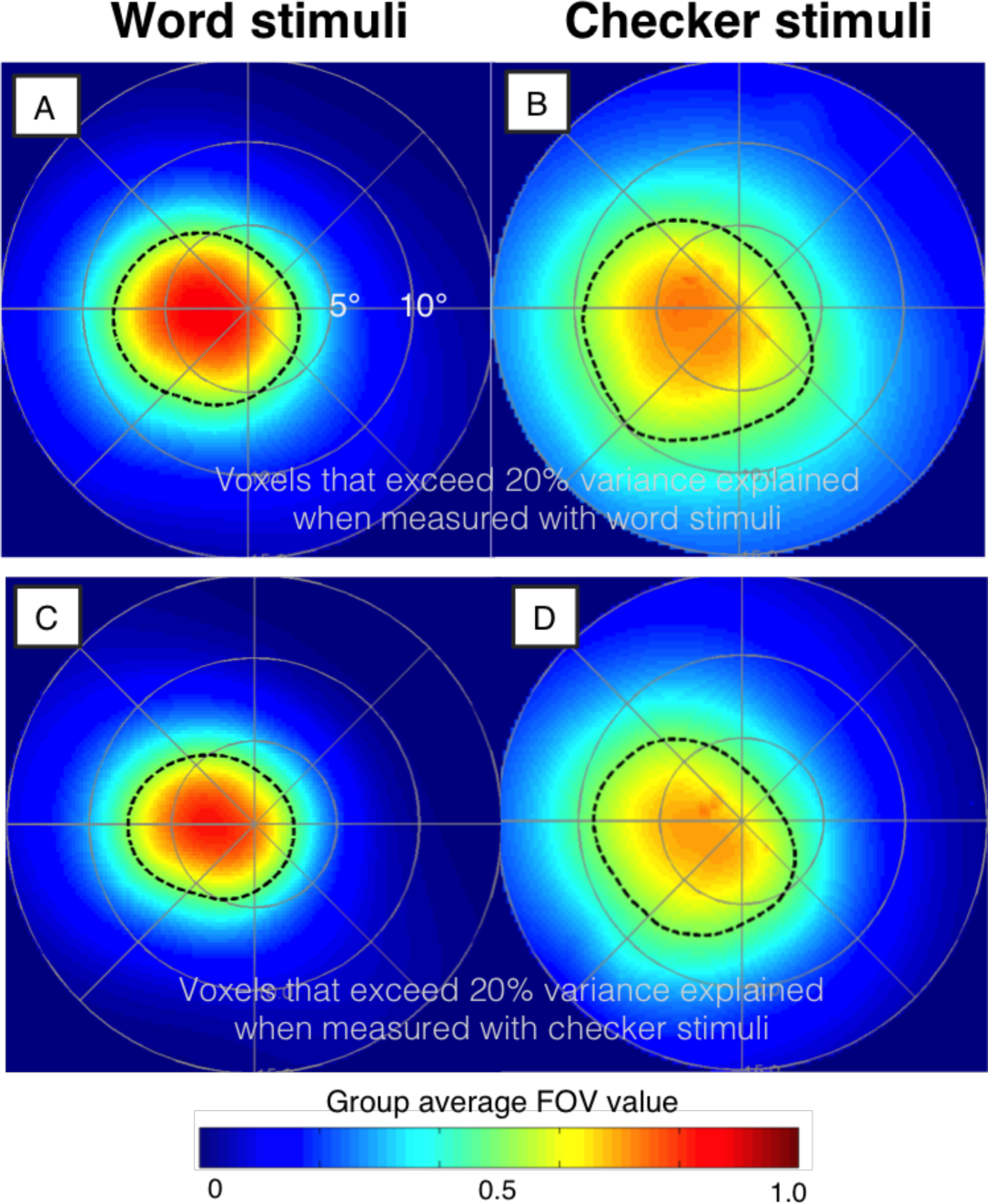
Group average right VOTRC FOVs for various subsets of voxels. Details as in Supplementary 4.

